# Müllerian-inhibiting substance (MIS) is both necessary and sufficient for testicular differentiation in Chinese soft-shelled turtle *Pelodiscus sinensis*

**DOI:** 10.1101/708073

**Authors:** Yingjie Zhou, Wei Sun, Han Cai, Haisheng Bao, Yu Zhang, Guoying Qian, Chutian Ge

## Abstract

Müllerian-inhibiting substance (*Mis*, or anti-müllerian hormone, *Amh*), a member of TGF-β superfamily, as initiator or key regulator in sexual development has been well documented in some vertebrates, especially in fish. However, its functional role has not been identified yet in reptiles. Here we characterized the *Mis* gene in Chinese soft-shelled turtle *Pelodiscus sinensis (P. sinensis)*, a typical reptilian species exhibiting ZZ/ZW sex chromosomes. The mRNA of *Mis* was initially expressed in male embryonic gonads by stage 15, preceding gonadal sex differentiation, and exhibited male-specific expression pattern throughout embryogenesis. Moreover, *Mis* was rapidly up-regulated during female-to-male sex reversal induced by aromatase inhibitor letrozole. Most importantly, *Mis* loss of function by RNA interference led to complete feminization of genetic male (ZZ) gonads, suppression of the testicular marker *Sox9*, and upregulation of the ovarian regulator *Cyp19a1*. Conversely, overexpression of *Mis* in ZW embryos resulted in female-to-male sex reversal, characterized by the formation of testis structure, ectopic activation of *Sox9*, and a remarkable decline in *Cyp19a1*. Collectively, these findings provide the first solid evidence that *Mis* is both necessary and sufficient to drive testicular development in a reptilian species, *P. sinensis*, highlighting the significance of the TGF-β pathway in reptilian sex determination.

## INTRODUCTION

In vertebrates, sex determination and gonadal differentiation generally follows the orderly expression of a series of sex-specific genes, which is triggered by primary sex-determining signal. Since the initial discovery of *Sry* in eutherian mammal (Sinclair *et al*. 1990; Koopman *et al*. 1990; Koopman *et al*. 1993), several sex-determining genes have been identified in some vertebrate species, such as *Dmrt1* in chicken (Smith *et al* 2009; Lambeth *et al* 2014), *Dmw* in frog (Yoshimoto *et al* 2008), *Foxl2* in goat(Boulanger *et al* 2014), *Dmy* (Matsuda *et al* 2002; Nanda *et al* 2002), *Amhr2* (Kamiya *et al* 2012), *SdY* (Yano *et al* 2012), *Gsdf* (Myosho *et al* 2012), *Sox3* (Takehana *et al* 2014), *Gdf6Y* (Reichwald *et al* 2015), *Amhy* (Hattori *et al* 2012; Li *et al* 2015) and *Dmrt1* (Chen *et al* 2014) in fish. Among these genes, *Amhy, Amhr2* and *Gsdf* are from the transforming growth factor beta (TGF-β) signaling pathway, suggesting a conserved role of this pathway in the primary sex determination in fish. However, whether the TGF-β pathway play a critical role in reptilian sex determination and differentiation has not yet been reported.

Müllerian inhibiting substance (*Mis*), also known as Anti-müllerian hormone (*Amh*), is a hormone-related gene belonging to TGF-β superfamily. *Mis* gene has been found and cloned in various vertebrates of different evolutionary positions, such as mouse (King *et al* 1991), chicken (Neeper *et al* 1996), American alligator (Western *et al* 1999), medaka (Klüver *et al* 2007), and tilapia (Shirak *et al* 2006). It functions through binding with the type II receptor *(AmhrII)*, which in turn induces the formation of receptor polymers to activate downstream target genes (Josso *et al* 2001; Rey *et al* 2003; Johnson *et al* 2008). In mammals, *Mis* gene is expressed in Sertoli cells of embryonic testes, and responsible for the regression of the Müllerian ducts, but it is not detected during the female embryonic development (Josso *et al* 2001). Like mammals, chicken *Mis* is expressed only in males and induce the regression of two Müllerian ducts (Smith *et al* 1999), however, knockdown of *Mis* in chicken ZZ embryos doesn’t alter gonadal development (Lambeth *et al* 2015). Despite the lack of Müllerian ducts in most teleost fish, the sexually dimorphic expression pattern of *Mis* and *AmhrII* is also detected in developing or mature gonads (Miura *et al* 2002; Yoshinaga *et al* 2004; Wu *et al* 2010; Eshel *et al* 2014). Deletion of *Amhy* in Patagonian pejerrey and *Amhr2* in *Takifugu rubripes*, both residing on Y sex chromosome, results in male-to-female sex reversal, thus rendering these two genes as male sex-determining genes (Kamiya *et al* 2012; Hattori *et al* 2012). Correlative studies in reptiles show that *Mis* exhibits male-specific embryonic expression, preceding the gonadal sex differentiation, in the red-eared slider turtle (Shoemaker *et al* 2007), painted turtle (Radhakrishnan *et al* 2017) and American alligator (Western *et al* 1999). These observations suggest a possible upstream position of *Mis* in the male pathway of reptiles, and its functional role in determining the gonadal sexual fate needs to be elucidated.

Chinese soft-shelled turtle *Pelodiscus sinensis (P. sinensis)* exhibiting ZZ/ZW genetic sex-determining system has been recently emerged as an ideal turtle model for investigating reptilian sex determination and differentiation, due to the well-established genetic modulation technique (Sun *et al* 2017; Ge *et al* 2017) and available genome resource (Wang *et al* 2013). In this study, we found that knockdown of *Mis* by RNA interference resulted in male-to-female sex reversal in *P. sinense*. Conversely, overexpression of *Mis* led to complete masculinization of female genetic turtles, indicating a both necessary and sufficient role of *Mis* to drive testicular development in a reptilian species.

## MATERIALS AND METHODS

### Eggs Incubation and Tissue Collection

Freshly laid Chinese soft-shelled turtle *(P. sinensis)* eggs were obtained from the Dafan turtle farm (Zhejiang, China). Fertilized eggs were placed in egg incubators at 31°C, with humidity maintained at 75%-85%. During the incubation process, embryos of different developmental stages, which were identified according to criteria established by Tokita and Kuratani (Tokita *et al* 2001), were removed from eggshells, decapitated and placed in PBS for gonad-mesonephros complexes (GMCs) and whole-gonads collection. GMCs were fixed in 4% paraformaldehyde (PFA) overnight at 4°C, dehydrated through 50% ethanol, and then stored in 70% ethanol at 4°C until paraffin embedding and sectioning was performed. Gonads were broken up thoroughly and immersed in TRIzol reagent (Invitrogen, USA) for total RNA isolation. Meanwhile, all embryos from treated and control groups were treated by liquid nitrogen grinding and then stored at −80°C for genomic DNA extraction. Additionally, adult turtle testis was prepared and stored at −80°C for *Mis* cDNA cloning. All animal experiments were carried out according to a protocol approved by Zhejiang Wanli University.

### Cloning of *P. sinensis Mis* cDNA

The total RNA from testis of adult turtle *P. sinensis* was extracted using TRIzol reagent (Invitrogen, USA). The first complementary DNA (cDNA) was then synthesized from 2μg of RNA by using the RevertAid™ First Strand cDNA Synthesis Kit (Fermentas, USA) following the manufacturer’s instructions. 5’ and 3’ RACE was carried out according to the manufacturer’s protocol of SMART RACE cDNA Amplification kit (Clontech, Takara, Japan). The sequences of primers for RACE are as follows: Mis-GSPF1: 5’-CGCTCTCCACCCGCATCCCCGACT-3’; *Mis-*GSPF2: 5’-GGTTTCTGCCTCGCTCTTCAGTCCT-3’; *Mis*-GSPR1: 5’-TACTGCAAAGCGACTC CTAGCAC-3’; *Mis*-GSPR2: 5’-TGGCAGACATTTCTCTTAGGGCTT-3’. The PCR products were extracted from agarose gel using MiniBEST Agarose Gel DNA Extraction Kit (Takara, Japan) based on manufacturer’s instructions, and cloned into pMD18-T (Takara) vector and then transformed into *E.coli* DH5α for sequencing. Alignment of deducted amino acid sequences were carried out by Clustal X software, and the phylogenetic tree was constructed using the Neighbour-Joining(N-J) method in Mega 6.0 software. The sequences of amino acid used in the phylogenetic analysis were obtained from GenBank (NCBI).

### Aromatase Inhibitor letrozole treatment

A non-steroidal aromatase inhibitor letrozole (Sigma, USA) were administered to eggs at developmental stage 15 and 16 (gonadal differentiation normally begins from late stage 17). The letrozole was dissolved in 95% ethanol at a concentration of 20 μg/μl, and 10 μl of drug was topically applied to the eggshell in the region adjacent to the embryo. Controls were treated with 10 μl of 95% ethanol. Gonad-mesonephros complexes were dissected from treated and control embryos at stage 27 for histology and immunohistochemistry. Gonads were separated from adjacent mesonephros at stage 17, 21 and 25, and stored for qRT-PCR analysis.

### Construction of LV-*Mis*-shRNA Vector System

The lentivirus vector was used to deliver shRNAs specifically targeting *Mis* mRNA into living embryos of Chinese soft-shell turtle before sexual differentiation, to knockdown endogenous *Mis* transcripts. The designed shRNA construct contained a unique 21 nt double-stranded *Mis* sequence that presented as an inverted complementary repeat, a loop sequence (5’-CTCGAG-3’) and the RNA Plo-II terminator (5’-TTTTTT-3’). Annealed oligonucleotides were ligated into pGP-U6 (GenePharma, Shanghai, China) between the *Bbs* and *Xho* sites by T4 DNA ligase (TaKaRa) to produce pGP-U6-*Mis*-shRNA. The pGP-U6-*Mis*-shRNA construct was digested with *Age*l*-EcoR*I and inserted into the *EcoR*I site of pGLV-U6-GFP (GenePharma). The lentivirus vector can also express green fluorescent protein (GFP), providing rapid visual assessment of the viral infection efficiency of embryos. The recombinant vector pGLV-GFP-*Mis*-shRNA was termed as LV-*Mis*-shRNA. The negative control vector (pGLV-GFP-NC-shRNA, termed as LV-NC-shRNA) contained a nonsense shRNA insert in order to control any effects caused by non-RNAi mechanisms. The sequence of the shRNA are as follows: *Mis*-shRNA (5’-GGTGCTGCATCTTGAGGAAGT-3’).

For the generation of lentivirus, 293T producer cells were transfected with optimized packaging plasmids (pGag/Pol, pRev and pVSV-G) along with pGLV-*Mis*-shRNA or pGLV-NC-shRNA expression clone constructs by lipofectamine. 24 h post transfection, the transfection mix was replaced by a fresh culture medium (without antibiotics). The virus-containing supernatant was harvested 72 h post transfection, cleared by centrifugation (3000 rpm/min, 15 min, and 4°C), and then filtered through a 0.45 μm filter (Millipore, USA). Viruses were titrated by adding serial dilutions to fresh 293 T and assessed using GFP expression after 48 h. Viral titres of approximately 1×10^9^ infectious units/ml were obtained. Lentivirus aliquots were stored at −80 °C before infection of turtle embryos.

### Construction of LV-*Mis*-OE Vector System

Total RNA was isolated from testis of adult Chinese soft-shelled turtle and then reverse transcription was performed to prepare the cDNA. The full-length open reading frame (1401bp) of *P. sinensis Mis* gene was PCR amplified from cDNA using forward primer 5’-CCCCAAATTGTAGAGGCGAACC-3’ and reverse primer 5’-TGAGGGCAGGGCAGAGGAGG-3’. The PCR product was digested with *EcoR*I and cloned to pGLV-EF1a-GFP (LV-4, GenePharma). The recombinant vector pGLV-GFP-*Mis* was named LV-*Mis*. The empty vector pGLV-GFP-empty was constructed as a negative control (LV-empty). High quality proviral DNA was used to transfect 293T cells. Virus propagation was carried out as described above.

### Infection of Turtle Embryos

A high-titre virus of LV-*Mis*-shRNA or LV-*Mis*-OE (at least 1×10^8^ infectious units/ml, 5 μl per embryo) was injected into turtle embryos at stage 14 before the time point (stage 15) that *Mis* began to exhibit a highly male(ZZ)-specific expression pattern, using a fine metal Hamilton needle (diameter: 0.5 mm). Each 200 eggs were injected in two treated groups, and 200 control eggs were injected with scrambled control virus of LV-NC-shRNA or LV-empty. Eggs were sealed with parafilm and incubated for the indicated time points (stage 25 and 27). Embryos showing robust GFP fluorescence in the urogenital system were chosen for further analysis.

### Embryo sexing

The genomic DNA was extracted from all tested embryos, and amplification of sex chromosome-specific DNA fragment was subsequently performed to identify the genetic sex of each embryo, which was well documented previously (Literman *et al* 2017). PCR products were visualized on 1% agarose gels. The lower bands represent Z-linked amplified fragments, and higher bands represent W-linked sex-diagnostic fragments (Fig.S3). The primer sequences for PCR are as follows: *Setd1b* (F: 5’-GATCGAATTACATCCTGC CT-3’, R:5’-TAAATTAG GACTGGAAGACACC-3’).

### Quantitative RT-PCR

Total RNA was extracted from embryonic gonads of different developmental stages, and subsequently synthesized for cDNA (methodology found above). Quantification of gene transcript levels in embryonic gonads of all treated and control groups was measured by qRT-RCR. In all PCR reactions, *Gapdh* was used as a reference gene. The qRT-RCR reaction was carried out using SYBR^®^ PrimeScript ^™^ II (Takara) in a Bio-Rad iCycler system. After normalization with *Gapdh*, relative RNA levels in samples were calculated using the comparative threshold cycle (Ct) method. Each RNA sample was analyzed in triplicate determinations. The primers sequences for PCR are as follows: *Gapdh* (F: 5’-GGC TTT CCG TGT TCC AAC TC-3’, R:5’-GAC AAC CTG GTC CTC CGT GTA TC-3’); Mis(F:5’-CGG CTA CTC CTC CCA CAC G-3’, R:5’-CCT GGC TGG AGT ATT TGA CGG-3’); *Cyp19*α*1*(F:5’-TCG TGG CTG TAC AAG AAA TAC GAA-3’, R:5’-CCA GTC ATA TCT CCA CGG CTC T-3’); *Sox9*(F:5’-TTT CCG ACC GCT AAA ACG ACA C-3’, R:5’-CTC CGC TGA CCA AAA CTT AGC CC-3’).

### Immunofluorescence

Gonad-mesonephros complexes (GMCs) were fixed in 4% PFA overnight at 4°C, dehydrated in graded ethanol, then embedded in paraffin wax and sectioned. Paraffin sections (5-6 μm) were deparaffinized and rehydrated prior to immersion in 10 mM sodium citrate buffer for 20 min at a sub-boiling temperature (96-99°C) for antigen retrieval. After blocked for 1 h in blocking solution (10% Normal Donkey Serum, 3% BSA (albumin from bovine serum), and 0.3%Triton X-100) at room temperature, sections were covered with primary antibodies and incubated overnight at 4°C, followed by washing (three times, 10 min each time) in washing solution (1% Normal Donkey Serum, 3% BSA, 0.3% Triton X-100), secondary antibodies incubation (2 h, room temperature, dark environment) and washing (same as above). The primary antibodies used in this analysis included rabbit anti-MIS (1:200, produced privately through Sangon Biotech), rabbit anti-VASA (1:500, Abcam), rabbit anti-SOX9 (1:500, Millipore) and mouse anti-CTNNB1 (1:250, Sigma). Primary antibodies were detected using secondary antibodies AlexFluor 488 donkey anti-rabbit IgG or AlexFluor 594 donkey anti-rabbit IgG, AlexFluor 488 donkey anti-mouse IgG (1:250, Invitrogen). Nuclei were stained with DAPI (286 nmol/L, Sigma) and then washed with 0.01 mol/L PBS (three times, 5 min each time). Fluorescence signals were observed under a fluorescence microscope (Ti-E, nickon) or confocal microscope (A1 Plus, Nickon).

### Statistical Analyses

Each experiment was independently repeated at least three times. All data was expressed as the means ± S.D. and analyzed by One-Way Duncan test and ANOVA using the SPSS software. For all analyses, a P-value < 0.05 was regarded as statistically significant (*, *P*<0.05; **, *P*<0.01; ***, *P*<0.001).

## RESULTS

### Characterization of *Mis* gene in *P. sinensis*

The full-length coding sequence of *P. sinensis Mis* was obtained by 5’ and 3’ RACE. The complete cDNA sequence of *P. sinensis Mis* was 3232 base pairs (bp) (accession number KY964412), with a 997 bp 5’ untranslated region (UTR), an open reading frame (ORF) of 1401 bp, and an 834 bp 3’ UTR (Supplementary Fig. 1A). The deduced MIS protein comprised 466 amino acids, which includes two characteristic functional domains of the TGF-β superfamily: AMH-N and TGF-β domain with ten canonical cysteine residues. The amino acid sequence of *P. sinensis* MIS shared 47%, 21.06%, 19.25%, 32.22%, 18.55%, and 11.30% identity with that of the red-eared slider turtle *(Trachemys scripta)*, human *(Homo sapiens)*, mice *(Mus musculus)*, chicken *(Gallus gallus)*, frog *(Xenopus laevis)* and zebra fish *(Danio rerio)*, respectively (Supplementary Fig. 1B). The phylogenetic tree also showed that *P. sinensis* MIS was evolutionarily most closely related to the red-eared slider turtle, followed by chicken and mice, and distantly related to fish (Supplementary Fig. 1C).

### Sexually dimorphic expression of *Mis* in gonads of *P. sinensis*

To find out whether *Mis* is involved in testicular development in *P. sinensis*, we first analyzed the expression profile of *Mis* in embryonic gonads of both sexes at different developmental stages. RNA-seq showed that *Mis* transcripts were detected and already expressed highly in the male gonads as early as stage 15. It exhibited male-specific embryonic expression during the critical sex determination period (stage 15 to 19), with female gonads showing extremely low expression level (Fig. 1A). The sex-dependent expression was further confirmed by qRT-PCR (Fig. 1B). We also examined the cellular localization of MIS protein in embryonic gonads at stage 17, when the gonads were still morphologically undifferentiated and appeared identical between sexes. Immunofluorescence showed that MIS protein was robustly expressed in Sertoli cells of the medullary sex-cords in male embryonic gonads, whereas the expression signals were undetectable in female gonads (Fig. 1C).

**Figure 1.**
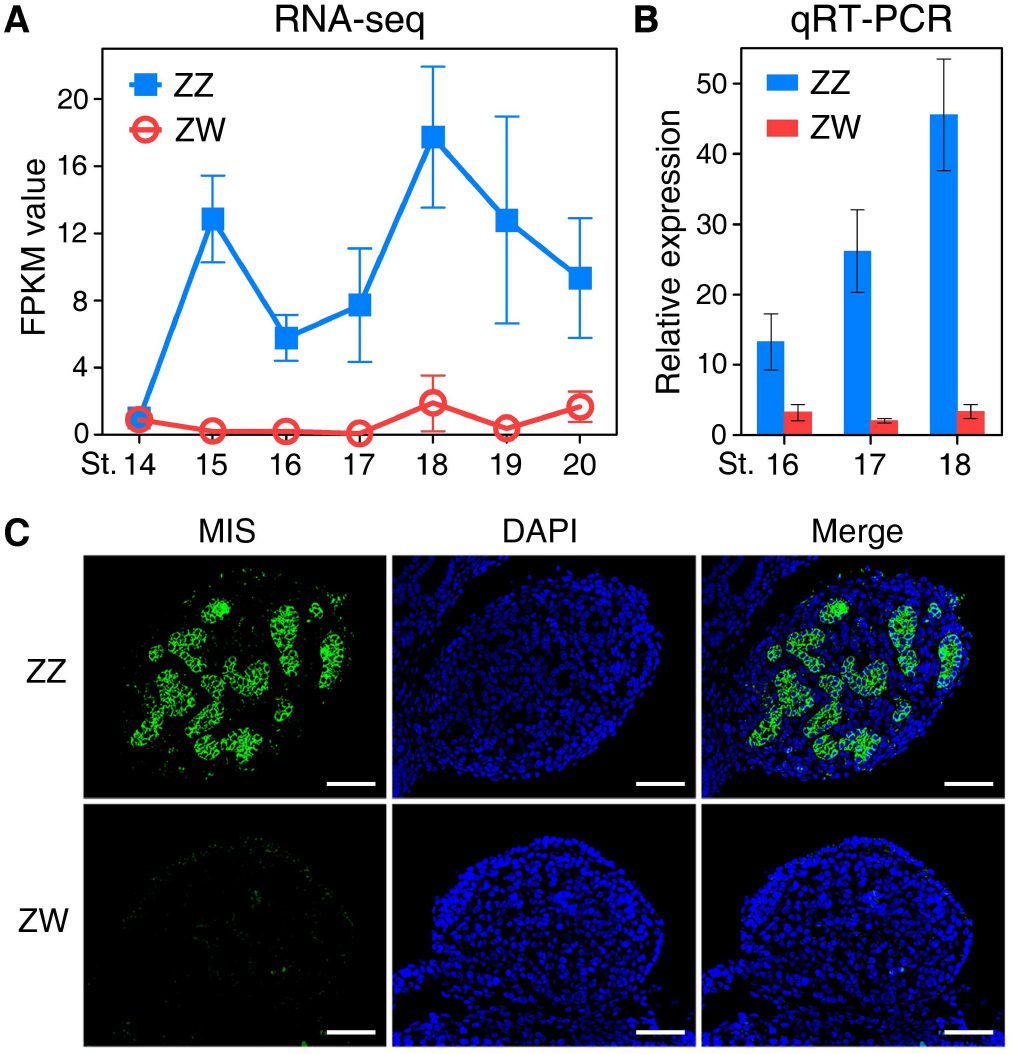
The sexually dimorphic expression of *Mis* in early embryonic gonads of *P. sinensis*. (A, B) The transcript expression levels of *Mis* in gonads of both sexes during the critical sex determination period (stage 15-19), determined by RNA-seq (A) and qRT-PCR (**B**). *Mis* exhibited a highly male-specific expression pattern in early embryonic gonads. Data are shown as means ± S.D. N≥3. (C) Immunofluorescence of MIS in male and female embryonic gonads at stage 17. MIS protein was robustly expressed in the medullary region of ZZ gonads. Scale bars are 50 μm.

### Upregulation of *Mis* in ZW gonads during female-to-male sex reversal

Treatment of aromatase inhibitor (AI) letrozole at early stages of sex determination (stage 15 and 16) induced ZW turtle embryos to develop towards the male phenotype (Fig. 2A). Tails of control ZW embryos were not beyond the hem of calipash, shorter than those in control ZZ embryos. However, tails of AI-treated ZW embryos became longer, with male genitals exposed from the cloacal orifice in most cases. The gonadal histological analysis showed that AI-treated ZW embryos exhibited medullary testis-cords and degenerated cortex (Fig. 2A). Furthermore, the testicular marker SOX9 was induced to be robustly expressed in medulla of the masculinized ZW gonads (Fig. 2B). These observations demonstrated that AI treatment at early stages indeed induced female-to-male sex reversal in *P. sinensis*.

**Figure 2.**
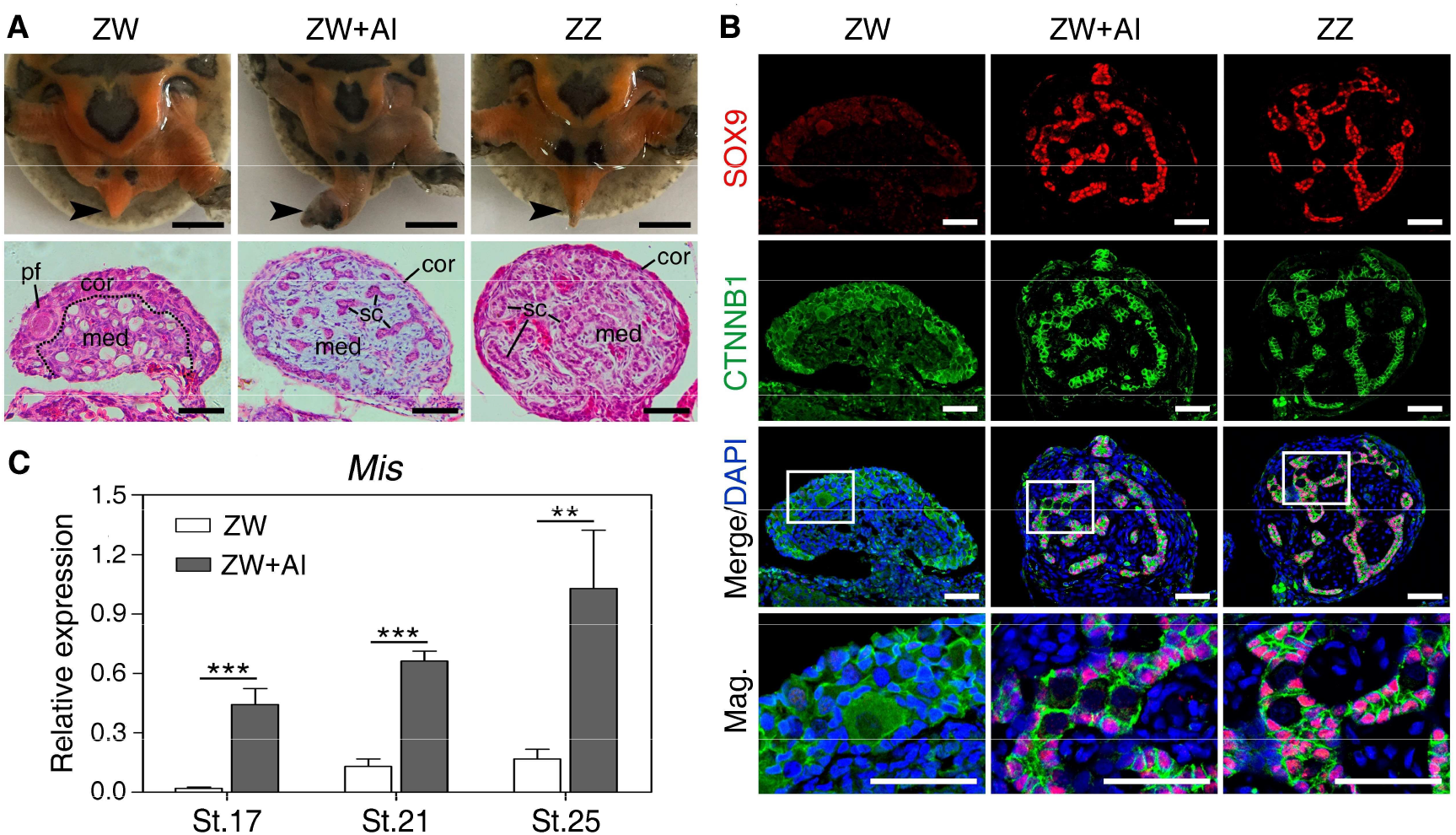
Upregulation of *Mis* in masculinized ZW gonads induced by aromatase inhibitor (AI). (A) Tail morphology (black arrow) and Hematoxylin and Eosin (H&E) staining of gonadal sections from control ZW, ZW+AI and control ZZ *P. sinensis* of stage 27. The male-to-female sex reversal were observed in ZW+AI gonads characterized by morphologically altered tail and medullary sex-cord formation. sc, sertoli cell; pf, primordial follicle; cor, cortical region; med, medullary region. Scale bars are 5 mm and 50 μm, respectively. (B) Double immunofluorescence of SOX9 and CTNNB1 in gonadal sections of control ZW, ZW+AI and control ZZ *P. sinensis* of stage 27. Ectopic expression of SOX9 protein were activated in masculinized medulla of ZW gonads. Scale bars are 50 μm. (C) The mRNA expression of *Mis* in ZW gonads with AI treatment at stage 17, 21 and 25, showing rapid and remarkable up-regulation, determined by qRT-PCR analysis. Data are shown as means ± S.D. N≥3. **, *P* < 0.01; ***, *P* < 0.001.

We next analyzed the expression changes of *Mis* in AI-induced female-to-male sex reversal to further determine whether *Mis* expression is associated with the testicular differentiation. qRT-PCR showed that *Mis* expression in ZW gonads increased dramatically in response to the female-to-male sex reversal (Fig. 2C). Intriguingly, the upregulation of *Mis* responded as early as stage 17, when the gonads were still morphologically undifferentiated between sexes, indicating that *Mis* is an early responder to the induction of male differentiation in *P. sinensis* (Fig. 2C).

### Feminization of ZZ turtle embryos with *Mis* knockdown

To investigate the function of *Mis* on male development of *P. sinensis*, we first established the *Mis* deficient turtle model by introducing shRNA against *Mis* in ovo at stage 14. qRT-PCR showed that the mRNA expression of Mis was >80% decreased in ZZ gonads from the embryos exhibiting global GFP reporter expression after LV-*Mis*-shRNA treatment than control ZZ gonads (LV-NC-shRNA) (Supplementary Fig. 2A, B). Phenotype of *Mis* deficient ZZ gonads were subsequently examined by gonadal histology and immunofluorescence. Control ZZ embryonic tails were straight and beyond the hem of calipash, but ZW embryonic tails were relative shorter and hid under the calipash (Fig. 3A, C). Control ZZ gonads were short and cylindrical, while ZW gonads were long and flat (Fig. 3D, F). In ZZ embryos with *Mis* knockdown, tails became curved and did not exceed the hem of calipash, and gonads became elongated and flat, exhibiting female-like morphology (Fig. 3B, E). Histological analysis of gonadal sections showed that the control ZZ gonads of stage 25 possessed a dense medulla with seminiferous cords and a degenerative cortex (Fig. 3G). Whereas control ZW gonads had a vacuolated medulla and a well-developed outer cortex (Fig. 3I). However, the *Mis* deficient ZZ gonads were completely feminized, characterized by a thickened cortex and a highly degenerated medulla (Fig. 3H). VASA staining showed that germ cells mainly located in medullary cords of control ZZ gonads, whereas control ZW gonads exhibited outer cortical distribution pattern of germ cells (Fig. 3J, L). VASA-positive germ cells in *Mis* deficient ZZ gonads displayed a female-like distribution, mainly enriched in the thickened cortex (Fig. 3K). Statistically, 32.8% (21 of 64) of genetic male embryos with *Mis* knockdown showed male-to-female sex reversal (Table 1).

**Table 1.** Phenotypes of embryos with knockdown or overexpression of *Mis*

| Viral treatment | No. of injected embryos | Embryos surviving until stage 25 | Genotype of embryos | Phenotype of embryos | sex reversal rate* |
| --- | --- | --- | --- | --- | --- |
| LV-NC-shRNA | 200 | 175 | ZZ:88; ZW:87 | M:88; F:87 | 0/88 |
| LV-*Mis*-shRNA | 200 | 138 | ZZ:64; ZW:74 | M:43; F:95 | 21/64 |
| LV-empty | 200 | 150 | ZZ:78; ZW:72 | M:78; F:72 | 0/72 |
| LV-*Mis*-OE | 200 | 124 | ZZ:62; ZW:62 | M:78; F:46 | 16/62 |

Genotype of embryos was identified by amplification of sex chromosome-specific DNA fragment.
Phenotype of embryos was assessed by gonadal histology, and SOX9 immunofluorescence.
\*Male-to-female sex reversal rate=No. of feminized genetic male embryos/total No. of ZZ embryos; female-to-male sex reversal rate=No. of masculinized genetic female embryos/total No. of ZW embryos.

**Figure 3.**
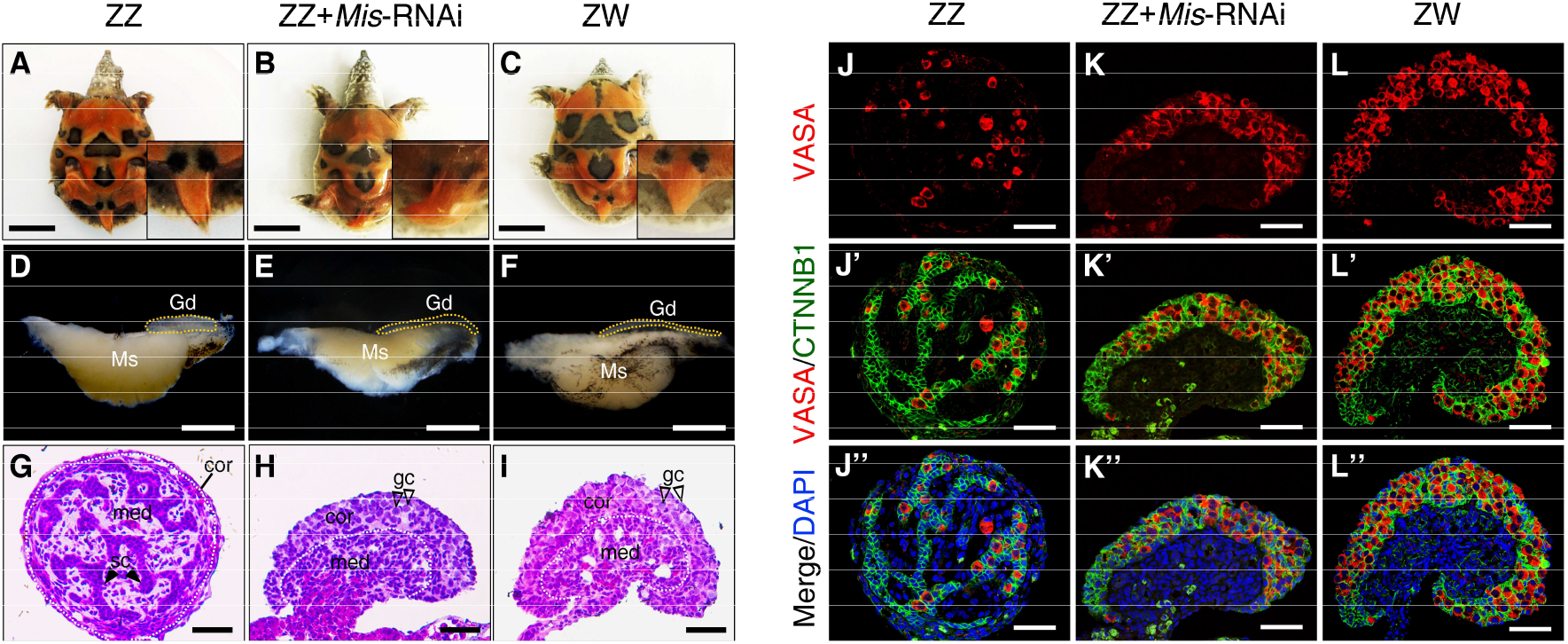
Feminization of ZZ embryos following Mis knockdown *in ovo*. (A-C) Morphology of tails from control ZZ, ZZ+Mis-RNAi and control ZW *P. sinensis* of stage 27. Scale bars are 1 cm. (D-F) Representative images of the gonad-mesonephros complexes (GMCs) from control ZZ, ZZ+ *Mis*-RNAi and control ZW embryos of stage 25. The ZZ gonads with *Mis* knockdown became elongated and flat, compared to control ZZ gonads. Gonads were outlined by yellow dotted lines. Gd, gonad; Ms, mesonephros. Scale bars are 1 mm. (G-I) H&E staining of gonadal sections from control ZZ, ZZ+Mis-RNAi and control ZW embryos of stage 25. The ZZ gonads with *Mis* knockdown appeared thickened outer cortex and degenerated testis cord in medullary region, similar to control ZW gonads. The white dotted lines showed the separation between cortical and medullar regions. sc, sertoli cell; gc, germ cells; cor, cortical region; med, medullary region. Scale bars are 50 μm. (J-L’’) VASA and CTNNB1 immunostaining of gonadal sections from control ZZ, ZZ+Mis-RNAi and control ZW embryos of stage 25. A female-typical distribution pattern of germ cells was observed in *Mis* deficient ZZ gonads. Scale bars are 50 μm.

To further confirm the activation of the female developmental pathway in *Mis* deficient ZZ embryos, we analyzed the expression changes of testicular differentiation marker *Sox9* and ovarian development regulator *Cyp19a1*. At the mRNA level, significant down-regulation of *Sox9*, and remarkable up-regulation of *Cyp19a1* were observed in ZZ gonads of stage 25 with *Mis* knockdown relative to controls (Fig. 4A, B). At the protein level, the expression signals of SOX9 was detected specifically in the nuclei of Sertoli cells in control ZZ gonads, but it was not observed in control ZW gonads. SOX9 expression in *Mis* deficient ZZ gonads was sharply reduced and almost disappeared (Fig. 4C). These results suggested that loss of *Mis* in ZZ turtle embryos led to male-to-female sex reversal.

**Figure 4.**
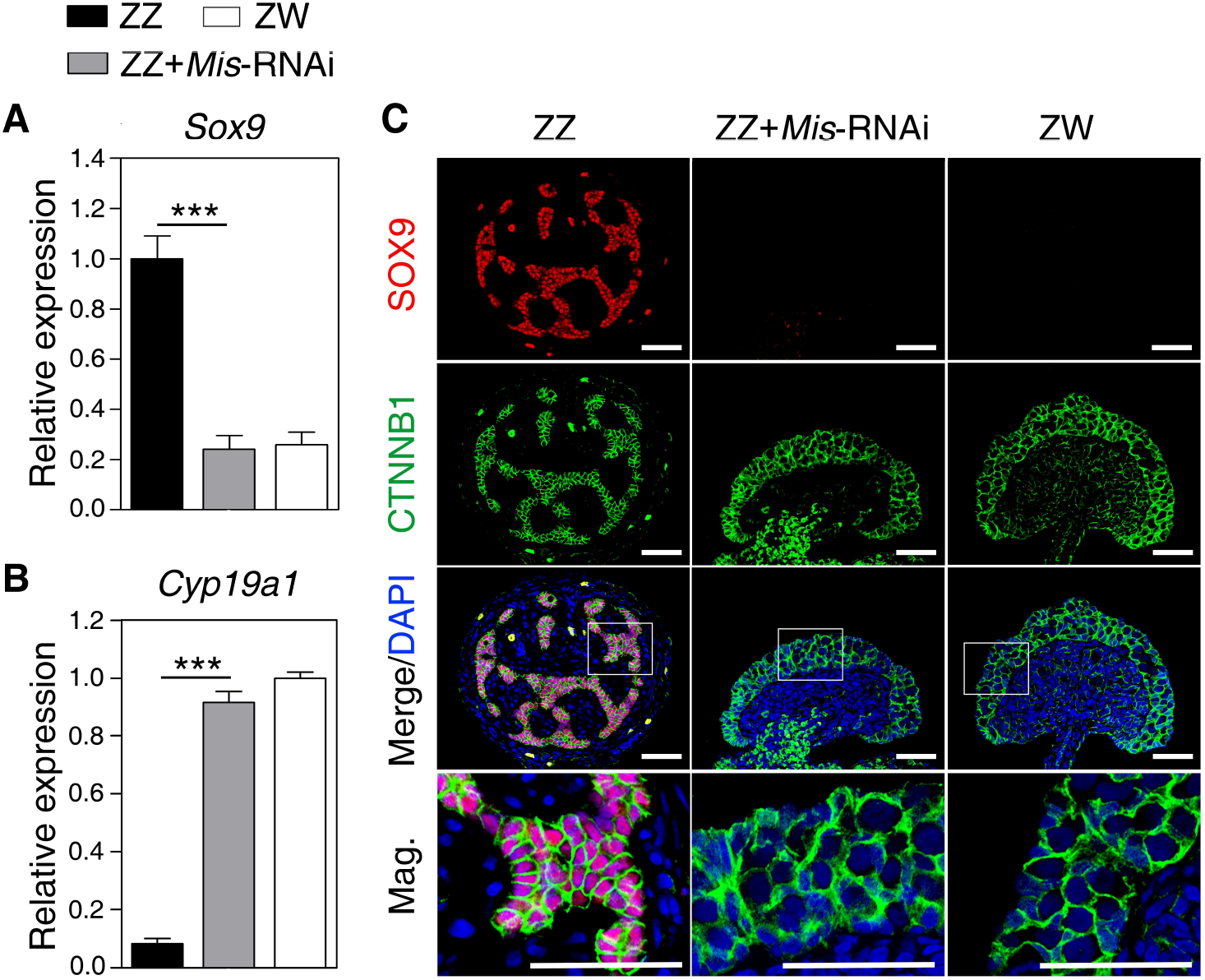
The *Sox9* and *Cyp19a1* expression change in response to Mis knockdown. (A, B) qRT-PCR of *Sox9* and *Cyp19a1* in control ZZ, ZZ+*Mis*-RNAi and control ZW gonads of stage 25, showing significantly reduced Sox9 expression and increased *Cyp19a1* expression in *Mis* deficient ZZ gonads. Data are shown as means ± S.D. N≥3. ***, P < 0.001. (C) Double immunofluorescence of SOX9 and CTNNB1 in sections of control ZZ, ZZ+*Mis*-RNAi and control ZW gonads of stage 25. SOX9 protein expression almost disappeared in Mis deficient ZZ gonads. Scale bars are 50 μm.

### Masculinization of ZW turtle embryos overexpressing *Mis*

The ectopic expression of *Mis* in ZW embryos was performed to determine if *Mis* was sufficient to initiate primary male differentiation in *P. sinensis*. *Mis*-overexpressing embryos were generated by injection of lentivirus vector carrying the *Mis* ORF into turtle eggs at stage 14 (Supplementary Fig. 2C). In ZW embryos overexpressing *Mis*, the tails became curved, and the gonads exhibited a short cylindrical structure, similar with control ZZ gonads (Fig. 5A-F). H&E staining of gonadal sections showed that ZW gonads overexpressing *Mis* exhibited a well-developed medulla with seminiferous cord-like structure (Fig. 5G-I). In LV-*Mis-OE* treated group, 25.8% (16 of 62) of ZW embryos showed female-to-male sex reversal (Table 1). Upregulation of *Sox9* and downregulation of *Cyp19a1* were observed in ZW gonads with *Mis* overexpression, determined by qRT-PCR (Fig. 6A, B). Ectopic activation of SOX9 protein in treated ZW gonads was further confirmed by immunofluorescence. Induced SOX9 expression was localized in the nuclei of Sertoli cells within the masculinized region (testis cords) in ZW gonads following *Mis* overexpression, but it seemed a little bit lower compared to control males (Fig. 6C). These data indicated that overexpression of *Mis* caused obvious masculinization of genetic female (ZW) embryos in *P. sinensis*.

**Figure 5.**
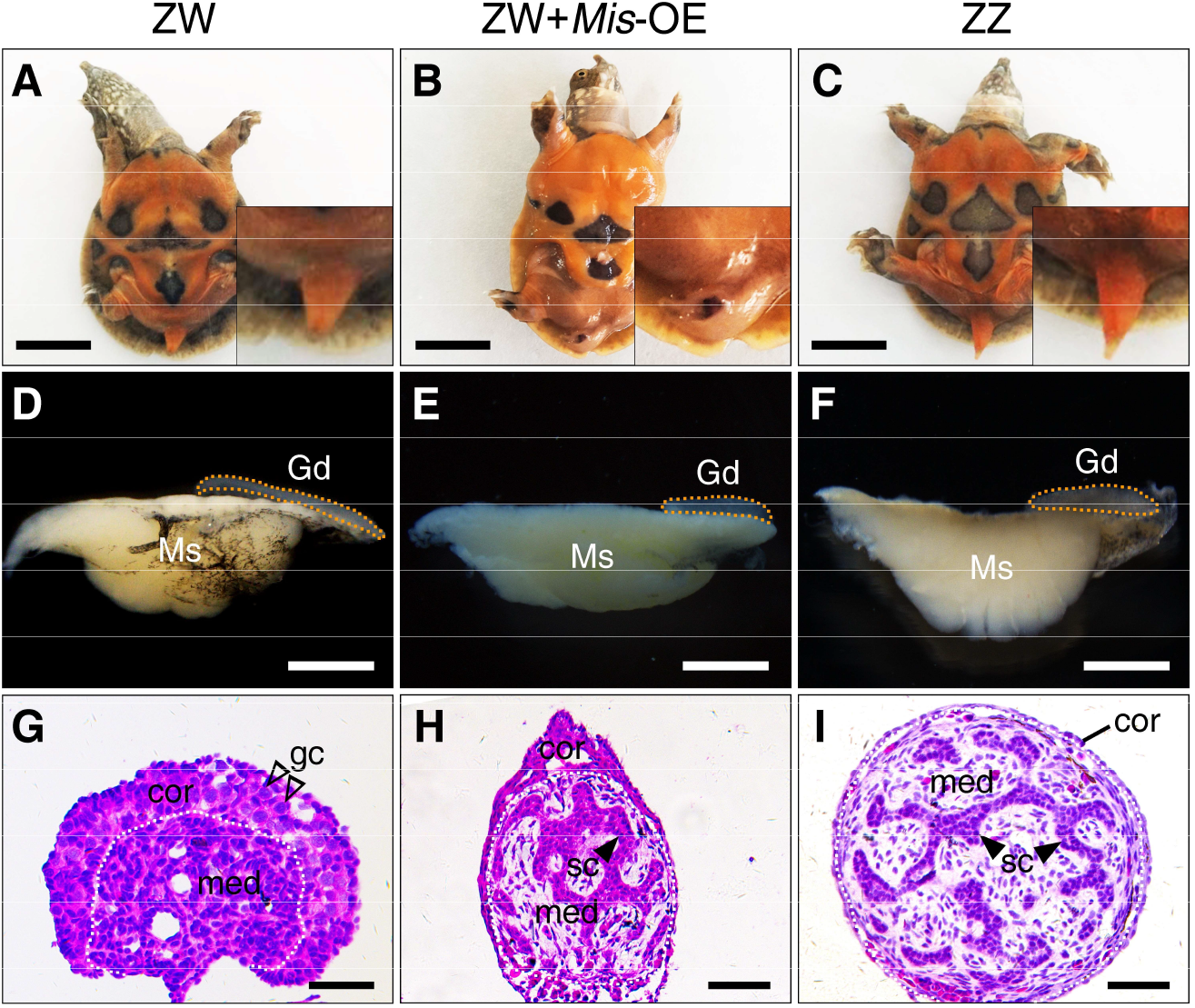
Masculinization of ZW embryos overexpressing *Mis in ovo*. The tails (A-C), GMCs (D-F) and H&E staining of gonadal sections (G-I) from control ZW, *ZW+Mis-OE* and control ZZ embryos. The ZW embryos overexpressing *Mis* showed the female-to-male sex reversal, characterized by curved tails and male-like gonads with seminiferous cord-like structure in medulla. Gd, gonad; Ms, mesonephros; sc, sertoli cell; gc, germ cells; cor, cortical region; med, medullary region. Scale bars are 1 cm (A-C), 1 mm (D-F) and 50 μm (G-I), respectively.

**Figure 6.**
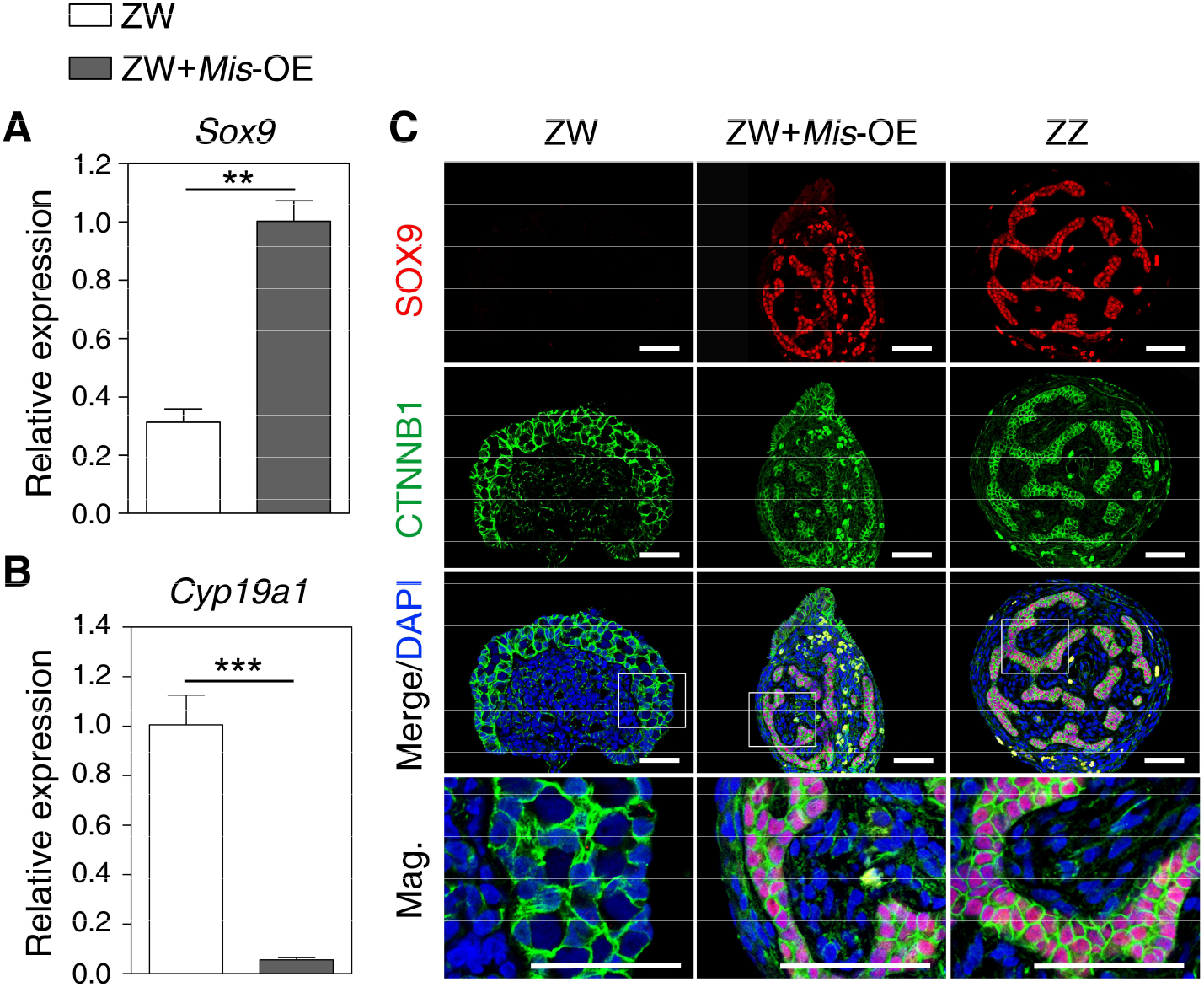
The *Sox9* and *Cyp19a1* expression change in response to Mis overexpression. (A, B) qRT-PCR of *Sox9* and *Cyp19a1* in control ZW, ZW+Mis-OE and control ZZ gonads of stage 25, showing increased Sox9 expression and reduced *Cyp19a1* expression in ZW gonads overexpressing *Mis*. Data are shown as means ± S.D. N≥3. **, P < 0.01; ***, *P* < 0.001. (C) Double Immunofluorescence of SOX9 and CTNNB1 in sections of control ZW, ZW+*Mis*-OE and control ZZ gonads of stage 25. SOX9 protein was induced to express robustly in gonadal medulla of ZW embryos overexpressing *Mis*. Scale bars are 50 μm.

## DISCUSSION

The conserved roles of TGF-β signaling pathway in sex determination have been recently functionally characterized in teleost fish, through the discoveries of three sex-determining genes, *Amhr2, Gsdf* and *Amhy* (Kamiya *et al* 2012; Myosho *et al* 2012; Hattori *et al* 2012). In this study, we provide the first solid evidence that *Mis* is both necessary and sufficient to induce male development in a reptilian species, *P*. *sinensis*, highlighting the significance of the TGF-β pathway in reptilian sex determination and sexual differentiation.

In this study, we found that the male gonad-specific expression of *P. sinensis Mis* has already appeared as early as stage 15, clearly preceding the onset of gonadal differentiation, indicating an upstream role of *Mis* in the male pathway of *P. sinensis*. This finding is consistent with previous studies in the red-eared slider turtle (Shoemaker *et al* 2007), painted turtle (Radhakrishnan *et al* 2017) and American alligator (Western *et al* 2012). In *T. scripta* with temperature-dependent sex determination, *Mis* expression in gonad was significantly higher at male-than female-producing temperature from stage 16 onwards, the beginning of temperature-sensitive sex determination period (Shoemaker *et al* 2007; Shoemaker-Daly *et al* 2010; Czerwinski *et al* 2016). These correlative studies strongly imply the conserved role of *Mis* in male development across reptilian species.

In non-mammalian vertebrates, estrogen and its synthetase aromatase play an important regulatory role in early gonadal sex differentiation. Exogenous estrogen and aromatase inhibitor (AI) can override the effects of primary sex-determination signals, including genetic and environmental factors, if applied during critical developmental periods (Crews 1994a; Crews 1994b; Smith *et al* 2003; Schulz *et al* 2007; Kobayashi *et al* 2008; Ge *et al* 2017;). Treatment of AI onto chicken ZW eggs was able to induce upregulation of *Dmrt1*, a Z chromosome-linked master sex-determining gene, ultimately resulting in female-to-male sex reversal (Smith *et al* 2003). It has been proposed that exogenous steroid hormones may redirect the differentiation direction of gonads by interacting with the sex-specific genes, especially those located on the upstream of sexual development pathway (Matsumoto *et al* 2012). In this study, *Mis* expression in *P*. *sinensis* ZW gonads responded rapidly to the AI-induced female-to-male sex reversal, prior to the sexual differentiation. The finding is consistent with the studies on zebrafish that reported the estrogen-induced alteration in *Mis* expression had already appeared at early stages of gonadal differentiation (Schulz *et al* 2007). These observations suggest that *Mis* is associated with testicular differentiation, and likely lies on the upstream of male pathway in *P. sinensis*.

To date, any member of TGF-β signaling pathway has not been functionally identified in reptiles, including turtles. Using an *in ovo* turtle gene-modulating approach developed previously (Sun *et al* 2017; Ge *et al* 2017), we found that knockdown of *Mis* led to complete feminization of genetic male (ZZ) gonads, including gonadal morphology and germ cell distribution pattern, as well as downregulation of testicular marker *Sox9* and upregulation of ovarian regulators *Cyp19a1*, indicating that *Mis* gene is essential for male gonadal differentiation in *P. sinensis*. This is similar to the functional roles of *Amhy*, the Y chromosome-linked duplicated copy of *Amh*, in two teleost fish (Hattori *et al* 2012; Li *et al* 2015). In Patagonian pejerrey, *Amhy* knockdown in XY embryos caused upregulation of *Cyp19a1a* and development of ovaries (Hattori *et al* 2012). Likewise, knockdown of *Amhy* in XY Nile Tilapia resulted in ovarian differentiation (Li *et al* 2015). Conversely, ectopic expression of *Mis* in *P*. *sinensis* ZW gonads induced the formation of sex cord-like structures with robust expression of SOX9 protein, implying that *Mis* is sufficient to initiate testicular differentiation in *P. sinensis*. As expected, the genetic female (XX) gonads overexpressing *Amhy* developed into testis in Nile Tilapia (Li *et al* 2015). Recently, we found the same loss-of- and gain-of-functional role of *Mis* in *T. scripta*, a turtle species with temperature-dependent sex determination (data not published), suggesting a conserved role for *Mis* in sex determination of turtle species, even with different sex determination systems. Our previous studies on *P. sinensis* have reported that the onset of *Mis* expression preceded *Sox9*, but later than *Dmrt1*, and *Dmrt1* overexpression caused an elevated expression of *Mis* and *Sox9* in ZW *P. sinensis* (Sun *et al* 2017). In this study, ectopic activation of *Sox9* occurred in response to *Mis* overexpression, which means that *Mis* could regulate *Sox9* in *P. sinensis. Mis* expression was also earlier than *Sox9* in chicken (Oreal *et al* 1998) and American alligators (Western *et al* 1999), however, the genetic position between *Mis* and *Sox9* was opposite in mammals. All these findings indicate that *P. sinensis Mis* acts as a positive regulator in the primary male sexual differentiation, and the network of *Dmrt1-Mis-Sox9* might be the effective component of testicular development in *P. sinensis*. Despite the necessary and sufficient role, *Mis* and *Dmrt1* seems not the master sex-determining gene, as both genes do not localize on the sex chromosome. Further investigation will be required to identify the master sex-determining gene in *P. sinensis*. Understanding the genetic link between the putative master gene and male or female effective components (such as *Mis)* may finally unravel the full mechanism of sex determination and differentiation in P. *sinensis*.

In conclusion, we demonstrate for the first time in reptiles that *Mis* is both necessary and sufficient to drive testicular development, thereby operating as an upstream regulator in the male pathway of Chinese soft-shelled turtle *Pelodiscus sinensis*. This study highlights a conserved role of a member of TGF-β signaling pathway, *Mis*, in reptilian sex determination and gonadal differentiation, and the direct upstream regulator of *Mis* needs to be identified.

## ACKNOWLEDGEMENTS

We thank Mr. Wei Song and Caisheng Wang for turtle eggs collection and incubation. This study was supported by the National Natural Science Foundation of China (31872960), National Key Research and Development Program (2018YFD0900203), Natural Science Foundation of Zhejiang Province for Distinguished Young Scholars (LR19C190001), the Basic Public Welfare Research Projects of Zhejiang Province (LGN19C190005), the Major Agricultural Project of Ningbo (2017C110012), the Zhejiang Provincial Project of Selective Breeding of Aquatic New Varieties (2016C02055-4), Zhejiang Provincial Top Key Discipline of Biological Engineering (KF2016005, ZS2018008). C.G. and G.Q. conceived and designed the study; Y.Z., W. S., H. C., H.B. and Y.Z. performed the experiments; Y.Z. and W.S. analyzed data; Y.Z., W.S. and C.G. co-wrote the manuscript. All authors read and approved the manuscript.

---

**Figure S1.**
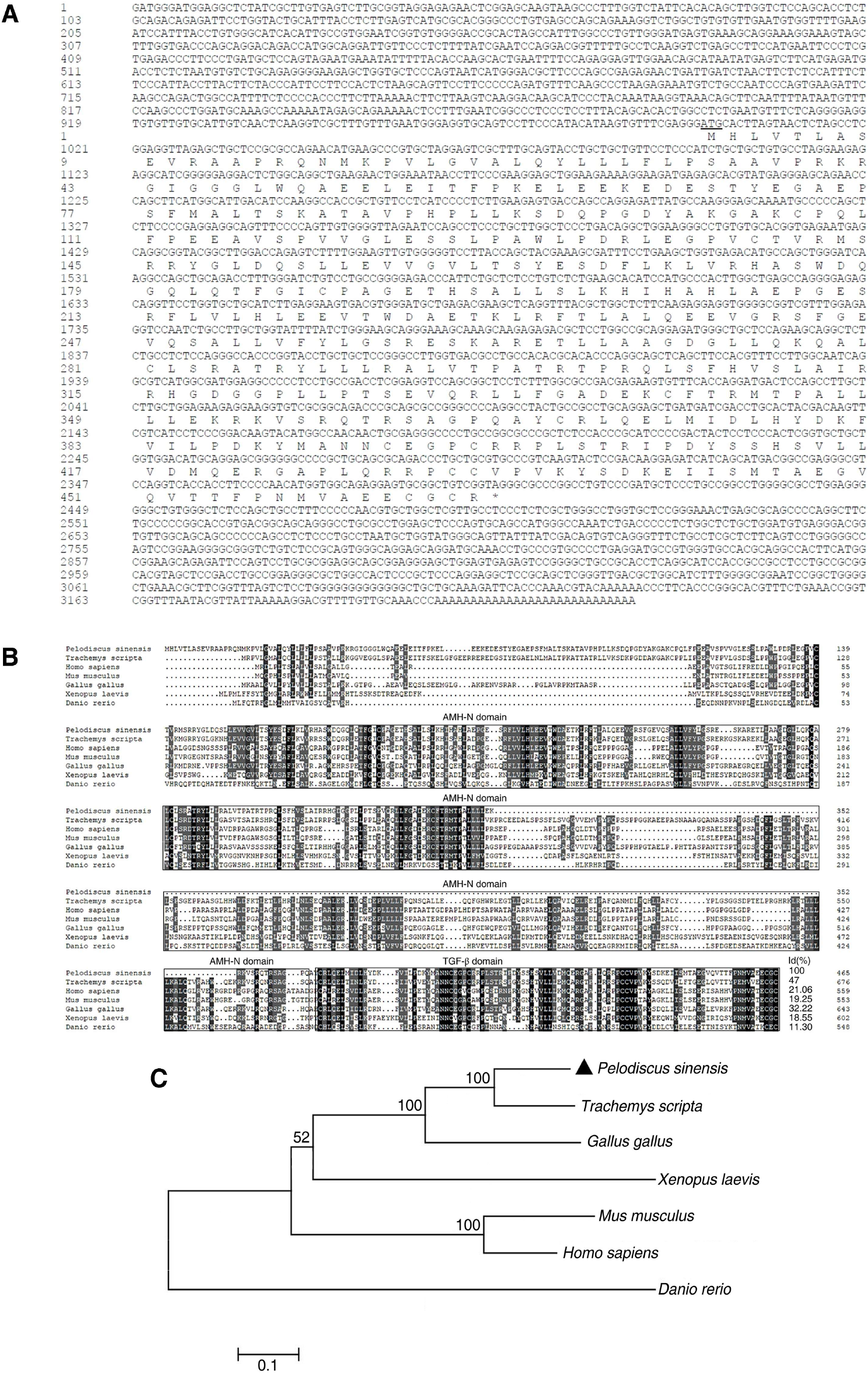
Sequence and phylogenetic analyses of *P. sinensis*. (A) The complete cDNA sequence of *P. sinensis Mis* and deduced amino acid sequence. The start codon ATG was underlined, and the stop codon was indicated by an asterisk. (B) Alignment of amino acid sequence of *P. sinensis* MIS with those from other typical species. The two characteristic functional domains of the TGF-β superfamily, AMH-N and TGF-β domain, were marked. (C) MIS phylogenetic tree from *P. sinensis* and other typical species based on Neighbor-Joining (N-J) method. Numbers at branches were confidence values based on 1000 bootstraps. Each branch length scale in terms of genetic distance was indicated above the tree.

**Figure S2.**
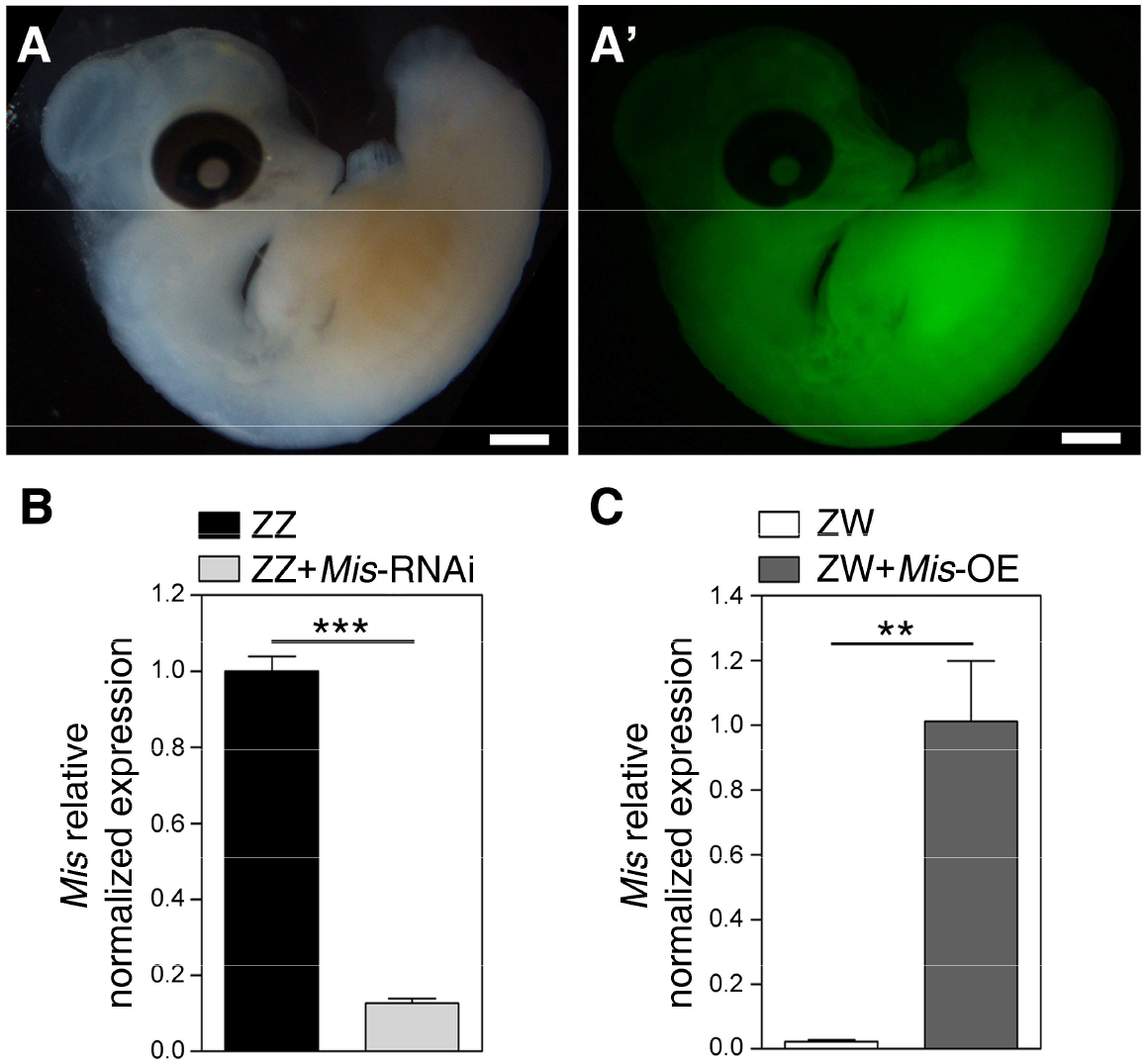
Establishment of *Mis*-Knockdown and -overexpressing turtle model using lentivirus vectors. (A, A’) The whole embryos of stage 15 infected with scrambled lentiviral vector (LV-NC) at stage 14 showed widespread GFP expression. Bright (A) and epifluorescence (A’) images. Scale bars are 1 mm. (B, C) qRT-PCR of Mis showed >80% downregulation in ZZ gonads with LV-*Mis*-shRNA treatment (B) and >50-fold upregulation in ZW gonads with *LV-Dmrt1-OE* treatment (C), respectively. Data are shown as means ± S.D. N≥3. **, P < 0.01; ***, *P* < 0.001.

**Figure S3.**
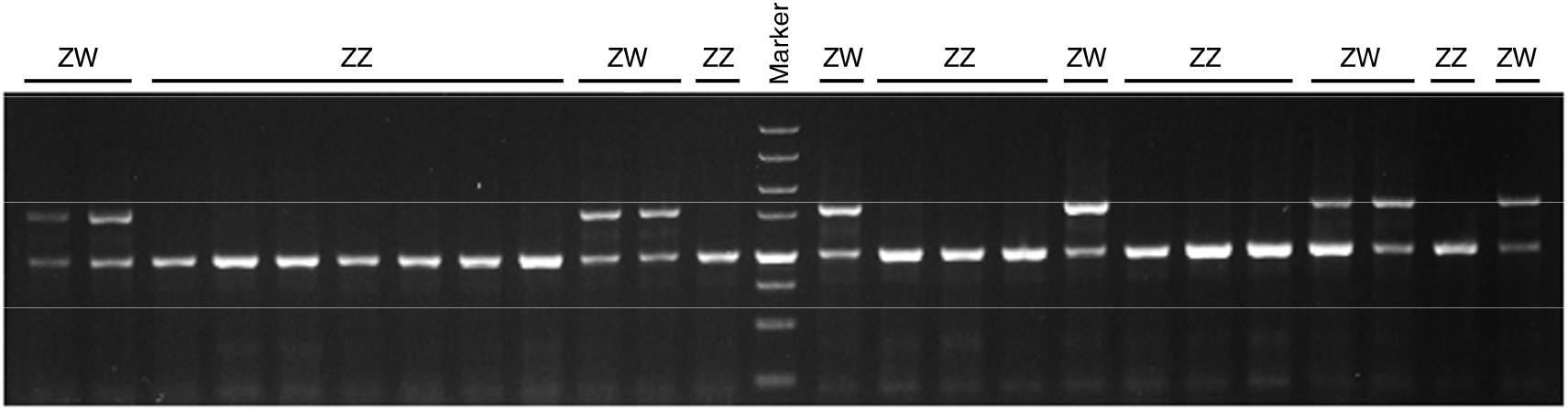
Sex-diagnostic amplification in *Pelodiscus sinensis*. Lower bands represent Z-linked amplified fragments, and higher bands represent W-linked sex-diagnostic fragments. One- and two-band indicated genetic male (ZZ) and female (ZW), respectively.

## LITERATURE CITED

Boulanger, L., M. Pannetier, L. Gall, A. Allais-Bonnet, M. Elzaiat, D. Le Bourhis et al., (2014) FOXL2 is a female sex-determining gene in the goat. Curr. Biol. 24: 404–408.

Chen, S., G. zhang, C. Shao, Q. Huang, G. Liu et al., (2014) Whole-genome sequence of a flatfish provides insights into ZW sex chromosome evolution and adaptation to a benthic lifestyle. Nat. Genet. 46: 253–260.

Czerwinski, M., A. Natarajan, L. Barske, L. L. Looger, B. Capel, (2016) A timecourse analysis of systemic and gonadal effects of temperature on sexual development of the red-eared slider turtle Trachemys scripta elegans. Dev. Biol. 420: 166–177.

Crews, D., (1994) Temperature, steroids and sex determination. J. Endocrinol. 142: 1–8.

Crews, D., J. M. Bergeron, (1994) Role of reductase and aromatase in sex determination in the red-eared slider (Trachemys scripta), a turtle with temperature-dependent sex determination. J. Endocrinol. 143: 279–289.

Eshel, O., A. Shirak, L. Dor, M. Band, T. Zak et al., (2014) Identification of male-specific Mis duplication, sexually differentially expressed genes and microRNAs at early embryonic development of Nile tilapia (Oreochromis niloticus). BMC Genomics 15: 774.

Ge, C., J. Ye, Y. Zhang, W. Sun, Y. Sang et al., (2017) Dmrt1 induces the male pathway in a turtle species with temperature-dependent sex determination. Development 144: 2222–2233.

Hattori, R. S., Y. Murai, M. Oura, S. Masuda, S. K. Majihi et al., (2012) A Y-linked anti-Mullerian hormone duplication takes over a critical role in sex determination. Proc. Natl. Acad. Sci. USA. 109: 2955–2959.

Josso, N., N. di, Clemente, L. Gouédard, (2001) Anti-Müllerian hormone and its receptors. Mol. Cell Endocrinol. 179: 25–32.

Johnson, P. A., T. R. Kent, M. E. Urick, and J. R. Giles, (2008) Expression and regulation of anti-Müllerian hormone in an oviparous species, the hen. Biol. Reprod. 78: 13–19.

Koopman, P., A. Münsterberg, B. Capel, N. Vivian, and R. Lovell-Badge, (1990) Expression of a candidate sex-determining gene during mouse testis differentiation. Nature 348: 450–452.

Koopman, P., J. Gubbay, N. Vivian, P. Goodfellow, and R. Lovell-Badge, (1991) Male development of chromosomally female mice transgenic for Sry. Nature 351: 117–21.

Kamiya, T., W. Kai, S. Tasumi, A. Oka, T. Matsunaga et al., (2012) A trans-species missense SNP in Misr2 is associated with sex determination in the tiger pufferfish, Takifugu rubripes (fugu). PLoS Genet. 8: e1002798.

King, T. R., B. K. Lee, R. R. Behringer, and E. M. Eicher, (1991) Mapping anti-Müllerian hormone (Mis) and related sequences in the mouse: identification of a new region of homology between MMU10 and HSA19p. Genomics 11: 273–283.

Kobayashi, T., H. Kajiura-Kobayashi, G. Guan, and Y. Nagahama, (2008) Sexual dimorphic expression of DMRT1 and Sox9a during gonadal differentiation and hormone-induced sex reversal in the teleost fish Nile tilapia (Oreochromis niloticus). Dev. Dyn. 237: 297–306.

Miura, T., C. Miura, Y. Konda, and K. Yamauchi, (2002) Spermatogenesis-preventing substance in Japanese eel. Development 129: 2689–2697.

Klüver, N., F. Pfennig, I. Pala, K. Storch, M. Schlieder et al., (2007) Differential expression of anti-Müllerian hormone (Mis) and anti-Müllerian hormone receptor type II (MisrII) in the teleost medaka. Dev. Dyn. 236: 271–281.

Lambeth, L. S., K. Ayers, A. D. Cutting, T. J. Doran, A. H. Sinclair et al., (2015) Anti-Müllerian hormone is required for chicken embryonic urogenital system growth but not sexual differentiation. Biol. Reprod. 93: 1–12.

Lambeth, L. S., C. S. Raymond, K. N. Roeszler, A. Kuroiwa, T. Nakata et al., (2014) Over-expression of DMRT1 induces the male pathway in embryonic chicken gonads. Dev. Biol. 389: 160–172.

Li, M., Y. Sun, J. Zhao, H. Shi, S. Zeng et al., (2015) A tandem duplicate of anti-Müllerian hormone with a missense SNP on the Y chromosome is essential for male sex determination in Nile Tilapia, Oreochromis niloticus. PLoS Genet. 11: e1005678.

Literman, R., S. Radhakrishnan, J. Tamplin, R. Burke, C. Dresser et al., (2017) Development of sexing primers in Glyptemys insculpta and Apalone spinifera turtles uncovers an XX/XY sex-determining system in the critically-endangered bog turtle Glyptemys muhlenbergii. Conserv. Genet. Resour. 9: 651–658.

Matsuda, M., Y. Nagahama, A. Shinomiya, T. Sato, C. Matsuda et al., (2002) DMY is a Y-specific DM-domain gene required for male development in the medaka fish. Nature 4l7: 559–563.

Myosho, T., H. Otake, H. Masuyama, M. Matsuda, Y. Kuroki et al., (2012) Tracing the emergence of a novel sex-determining gene in medaka, Oryzias luzonensis. Genetics 191: 163–170.

Matsumoto, Y., and D. Crews, (2012). Molecular mechanisms of temperature-dependent sex determination in the context of ecological developmental biology. Mol. Cell. Endocrinol. 354: 103–110.

Nanda, I., M. Kondo, U. Hornung, S. Asakawa, C. Winkler et al., (2002) A duplicated copy of DMRT1 in the sex-determining region of the Y chromosome of the medaka, Oryzias latipes. Proc. Natl. Acad. Sci. USA. 99: 11778–11783.

Neeper, M., R. Lowe, S. Galuska, K. J. Hofmann, R. G. Smith et al., (1996) Molecular cloning of an avian anti-Müllerian hormone homologue. Gene 176: 203–209.

Oreal, E., C. Pieau, M. G. Mattei, N. Josso, J. Y. Picard et al., (1998) Early expression of MIS in chicken embryonic gonads precedes testicular SOX9 expression. Dev. Dyn. 212: 522–532.

Reichwald, K., A. Petzold, P. Koch, B. R. Downie, N. Hartmann et al., (2015) Insights into Sex Chromosome Evolution and Aging from the Genome of a Short-Lived Fish. Cell 163: 1527–1538.

Rey, R., C. Lukas-Croisier, C. Lasala and P. Bedecarrás, (2003) MIS/MIS: what we know already about the gene, the protein and its regulation. Mol. Cell Endocrinol. 211: 21–31.

Radhakrishnan, S., R. Literman, J. Neuwald, A. Severin, and N. Valenzuela, (2017) Transcriptomic responses to environmental temperature by turtles with temperature-dependent and genotypic sex determination assessed by RNAseq inform the genetic architecture of embryonic gonadal development. PLOS One 12: e0172044.

Sinclair, A. H., P. Berta, M. S. Palmer, J. R. Hawkins, B. L. Griffiths et al., (1990) A gene from the human sex-determining region encodes a protein with homology to a conserved DNA-binding motif. Nature 346: 240–244.

Smith, C. A., M. Katz, and A. H. Sinclair, (2003) DMRT1 is upregulated in the gonads during female-to-male sex reversal in ZW chicken embryos. Biol. Reprod. 68: 560–570.

Smith, C. A., K. N. Roeszler, T. Ohnesorg, D. M. Cummins, P. G. Farlie, et al., (2009) The avian Z-linked gene DMRT1 is required for male sex determination in the chicken. Nature 461: 267–271.

Smith, C. A., M. J. Smith, and A. H. Sinclair, (1999) Gene expression during gonadogenesis in the chicken embryo. Gene 234: 395–402.

Shirak, A., E. Seroussi, A. Cnaani, A. E. Howe, R. Domokhovsky et al., (2006) Mis and Dmrta2 genes map to tilapia (Oreochromis spp.) linkage group 23 within quantitative trait locus regions for sex determination. Genetics 174: 1573–1581.

Shoemaker-Daly, C. M., K. Jackson, R. Yatsu, Y. Matsumoto, and D. Crews, (2010) Genetic network underlying temperature-dependent sex determination is endogenously regulated by temperature in isolated cultured Trachemys scripta gonads. Dev. Dyn. 239: 1061–1075.

Shoemaker, C., M. Ramsey, J. Queen, and D. Crews, (2007) Expression of Sox9, Mis and Dmrt1 in the gonad of a species with temperature-dependent sex determination. Dev. Dyn. 236: 1055–1063.

Sun, W., H. Cai, G. Zhang, H. Zhang, H. Bao et al., (2017) Dmrt1 is required for primary male sexual differentiation in Chinese soft-shelled turtle Pelodiscus sinensis. Sci. Rep. 7: 4433.

Schulz, R. W., J. Bogerd, R Male, J. Ball, M. Fenske et al., (2007) Estrogen induced alterations in Mis and Dmrt1 expression signal for disruption in male sexual development in the zebrafish. Environ. Sci. Technol. 41: 6305–6310.

Takehana, Y., M Matsuda, T. Myosho, M. L. Suster, K. Kawakami et al., (2014) Co-option of Sox3 as the male-determining factor on the Y chromosome in the fish Oryzias dancena. Nat. Commun. 5: 4157.

Tokita, M., and S. Kuratani, (2001) Normal embryonic stages of the Chinese softshelled turtle Pelodiscus sinensis (Trionychidae). Zool. Sci. 18: 705–715.

Western, P. S., J. L. Harry, J. A. Graves, and A. H. Sinclair, (1999) Temperature-dependent sex determination in the American alligator: MIS precedes SOX9 expression. Dev. Dyn. 216: 411–419.

Wu, G. C., P. C. Chiu, Y. S. Lyu, and C. F Chang, (2010) The expression of Mis and Misr2 is associated with the development of gonadal tissue and sex change in the protandrous black porgy, Acanthopagrus schlegeli. Biol. Reprod. 83: 443–453.

Wang, Z., J. Pascual-Anaya, A. Zadissa, W. Li, Y. Niimura et al., (2013) The draft genomes of soft-shell turtle and green sea turtle yield insights into the development and evolution of the turtle-specific body plan. Nat. Genet. 45: 701–706.

Yoshimoto, S., E. Okada, H. Umemoto, K. Tamura, Y. Uno et al., (2008) A W-linked DM-domain gene, DM-W, participates in primary ovary development in Xenopus laevis. Proc. Natl. Acad. Sci. USA. 105: 2469–2474.

Yano, A., R. Guyomard, B. Nicol, E. Jouanno, E. Quillet et al., (2012) An immune-related gene evolved into the master sex-determining gene in rainbow trout, Oncorhynchus mykiss. Curr. Biol. 22: 1423–1428.

Yoshinaga, N., E. Shiraishi, T. Yamamoto, T. Iguchi, S. Abe et al., (2004) Sexually dimorphic expression of a teleost homologue of Müllerian inhibiting substance during gonadal sex differentiation in Japanese flounder, Paralichthys olivaceus. Biochem. Biophys. Res. Commun. 322: 508–513.

